# A Motion Transformer for Single Particle Tracking in Fluorescence Microscopy Images

**DOI:** 10.1101/2023.07.20.549804

**Authors:** Yudong Zhang, Ge Yang

## Abstract

Single particle tracking is an important image analysis technique widely used in biomedical sciences to follow the movement of sub-cellular structures, which typically appear as individual particles in fluorescence microscopy images. In practice, the low signal-to-noise ratio (SNR) of fluorescence microscopy images as well as the high density and complex movement of subcellular structures pose substantial technical challenges for accurate and robust tracking. In this paper, we propose a novel Transformer-based single particle tracking method called Motion Transformer Tracker (MoTT). By using its attention mechanism to learn complex particle behaviors from past and hypothetical future tracklets (i.e., fragments of trajectories), MoTT estimates the matching probabilities between each live/established tracklet and its multiple hypothesis tracklets simultaneously, as well as the existence probability and position of each live tracklet. Global optimization is then used to find the overall best matching for all live tracklets. For those tracklets with high existence probabilities but missing detections due to e.g., low SNRs, MoTT utilizes its estimated particle positions to substitute for the missed detections, a strategy we refer to as relinking in this study. Experiments have confirmed that this strategy substantially alleviates the impact of missed detections and enhances the robustness of our tracking method. Overall, our method substantially outperforms competing state-of-the-art methods on the ISBI Particle Tracking Challenge datasets. It provides a powerful tool for studying the complex spatiotemporal behavior of subcellular structures. The source code is publicly available at https://github.com/imzhangyd/MoTT.git.

## 1 Introduction

A commonly used method to observe the dynamics of subcellular structures, such as microtubule tips, receptors, and vesicles, is to label them with fluorescent probes and then collect their videos using a fluorescence microscope. Since these subcellular structures are often smaller than the diffraction limit of visible light, they often appear as individual particles with Airy disk-like patterns in fluorescence microscopy images, as shown e.g., in Fig 1. To quantitatively study the dynamic behavior of these structures in live cells, these trajectories need to be recovered using single particle tracking techniques [14].

**Fig. 1.**
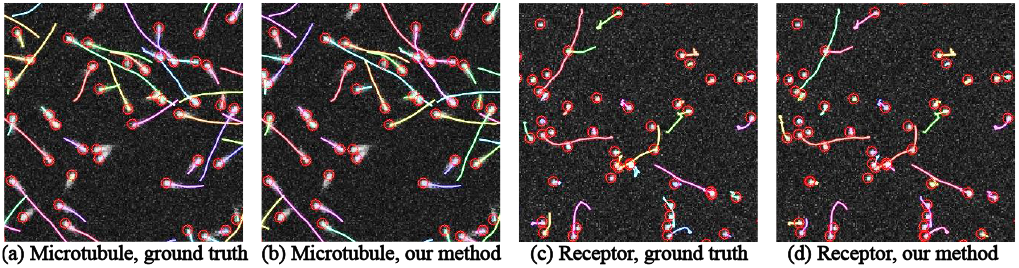
Tracking performance of our method. (a-b) ground truth trajectories of microtubule tips in (a) versus trajectories recovered by our method in (b). (c-d) ground truth trajectories of receptors in (c) versus trajectories recovered by our method in (d). (a-d) colors are chosen randomly to differentiate between individual trajectories.

Most single particle tracking methods follow a two-step paradigm: particle detection and particle linking. Specifically, particles are detected first in each frame of the image sequence. The detected particles are then linked between consecutive frames to recover their complete trajectories. The contributions of this paper focus on particle linking. Classical particle linking methods [5, 14, 9] are usually based on joint probability data association (JPDA) [10, 20], multiple hypothesis tracking (MHT) [19, 16], etc. Many classical methods have been developed and evaluated in the 2012 International Symposium on Biomedical Imaging (ISBI) Particle Tracking Challenge [6]. However, classical methods require manual tuning of many model parameters and are usually designed for a specific type of dynamics, making it difficult to apply to complex dynamics. In addition, the performance of these methods tends to degrade when tracking dense particles.

Deep learning provides a technique for automatically learning feature patterns and has been bringing performance improvements to many tasks. Recently, many deep learning-based single particle tracking methods have been developed. Many methods [30, 26, 25, 21] use long short-term memory (LSTM) [13] modules to learn particle behavior. However, in [30], the matching probabilities between each tracklet and its multiple candidates are calculated independently, and there is no information exchange between multiple candidates. In [30, 26], only detections in the next frame are used as candidates, which contain fewer motion features compared to hypothetical future tracklets. In [25, 21], the number of their subnetworks grows exponentially with the depth of the hypothesis tree, making the network huge. And the trajectories will be disconnected due to missing detections. In addition, the source codes of most deep learning-based single particle tracking methods are not available, making them difficult to use for non-experts.

Cell tracking is closely related to particle tracking. There are different classes of cell tracking methods. An important category is tracking-by-evolution [7], which assumes spatiotemporal overlap between corresponding cells. It is not suitable for tracking particle because they generally do not overlap between frames. Another important category is tracking-by-detection. Some methods [29, 18] in this category assume coherence in motion of adjacent cells, which is not suitable for tracking particles that move independently from each other. There are also cell tracking methods [2] that rely on appearance features, which are not suitable for tracking particles because they lack appearance features.

Transformer [27] is originally proposed for modeling word sequences in machine translation tasks and has been used in various applications [4, 3]. Recently, there have been many Transformer-based methods for motion forecasting [11, 17, 23], which improve the performance of motion forecasting in natural scenes (e.g., pedestrians, cars.). Compared to LSTM, Transformer shows advantages in sequence modeling by using the attention mechanism instead of sequence memory. However, to the best of our knowledge, Transformer has not been used for single particle tracking in fluorescence microscopy images.

In this paper, we propose a Transformer-based single particle tracking method MoTT, which is effective for different motion modes and different density levels of subcellular structures. The main contributions of our work are as follows: (1) We have developed a novel Transformer-based single particle tracking method MoTT. The attention mechanism of the Transformer is used to model complex particle behaviors from past and hypothetical future tracklets. To the best of our knowledge, we are the first to introduce Transformer-based networks to single particle tracking in fluorescence microscopy images; (2) We have designed an effective relinking strategy for those disconnected trajectories due to missed detections. Experiments have confirmed that the relinking strategy substantially alleviates the impact of missed detections and enhances the robustness of our tracking method; (3) Our method substantially outperforms competing state-of-the-art methods on the ISBI Particle Tracking Challenge dataset [6]. It provides a powerful tool for studying the complex spatiotemporal behavior of subcellular structures.

## 2 Method

Our particle tracking approach follows the two-step paradigm: particle detection and particle linking. We first use the detector DeepBlink [8] to detect particles at each frame. The detections of the first frame are initialized as the live tracklets. On each subsequent frame, we execute our particle linking method in four steps as follows. First (2.1), for each live tracklet, we construct a hypothesis tree to generate its multiple hypothesis tracklets. Second (2.2), all tracklets are pre-processed and then fed into the proposed MoTT network to predict matching probabilities between each live tracklet and its multiple hypothesis tracklets, as well as the existence probability and position of each live tracklet in the next frame. Third (2.3), we formulate a discrete optimization model to find the overall best matching for all live tracklets by maximizing the sum of the matching probabilities. Finally (2.4), we design a track management scheme for trajectory initialization, updating, termination, and relinking.

### 2.1 Hypothesis tree construction

Assuming that the particle linking has been processed up to frame *t*. In order to find correspondence between the current live tracklets and the detections of frame *t* + 1, we will build a hypothesis tree of depth *d* for each live tracklet, with its detection at frame *t* as the root node. To build the tree beyond the root node, we select *m* (real) detections of the next frame nearest to the current node as well as another null detection that represents a missing detection as children of the current node. If the current node is null, *m* (real) detections of the next frame nearest to the parent of the current node are selected. From the hypothesis tree, (*m* + 1)^*d*^ hypothesis tracklets will be obtained. Fig. 2 shows an example of the hypothesis tree construction with *m* = 2 and *d* = 2.

**Fig. 2.**
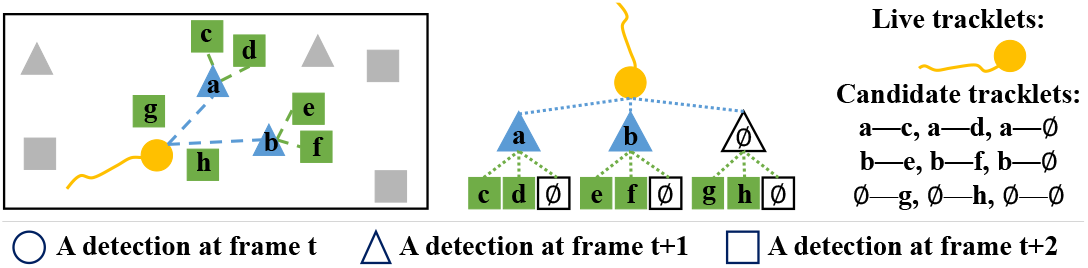
An example of hypothesis tree construction with *m* = 2 and *d* = 2

### 2.2 MoTT network

As shown in Fig. 3, We have designed a Transformer-based network, which contains a Transformer and two prediction head modules: classification head and regression head. Compared to the original Transformer, both the query masking and the positional encoding on the decoder are removed, since the input of the decoder is an unordered tracklet set. The classification head and regression head are constructed by fully connected layers.

**Fig. 3.**
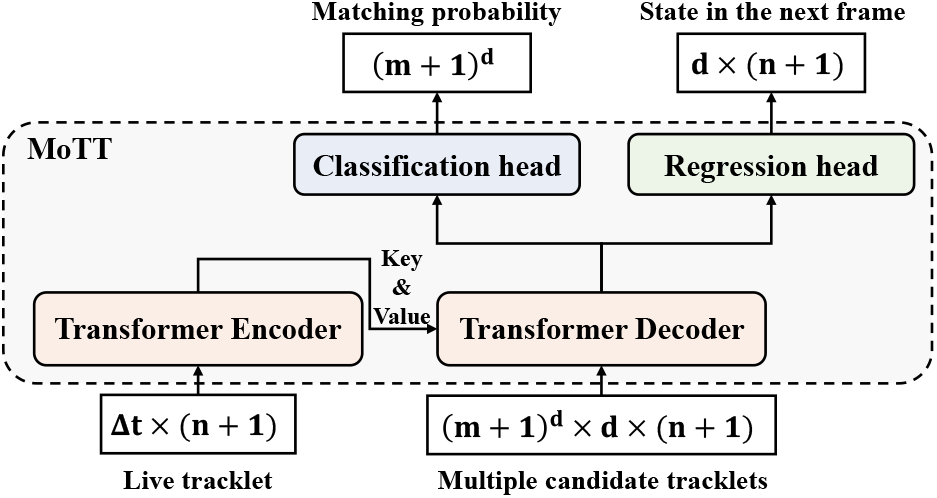
MoTT network structure. *Δt* is the constant length of live tracklets, *n* + 1 is the dimension number with the existence flag, *d* is the extended depth of hypothesis trees, *m* + 1 is the number of hypothesis tracklets. See supplementary material for the details of the MoTT structure.

For the generated tracklets from the previous step, the preprocessing is performed to make the length of all live tracklets equal to *Δt*, to convert position sequence to velocity sequence, and to add the existence flag making the coordinate dimension *n*+1. See supplementary material for the details of preprocessing. Then the preprocessed live tracklet is fed into the Transformer encoder, while the (*m* + 1)^*d*^ preprocessed hypothesis tracklets are fed into the Transformer decoder. The self-attention modules in the encoder and decoder are used to extract features of live tracklets and hypothesis tracklets, respectively. The cross-attention module is used to calculate the affinity between the live tracklet and its multiple candidate tracklets. The classification head outputs the predicted matching probabilities between the live tracklet and (*m* + 1)^*d*^ hypothesis tracklets. The regression head outputs the predicted existence probability and velocity of each live tracklet in the next frame. The existence probability represents the probability of the live tracklet existence in the next frame. The predicted velocity can be easily converted to the predicted position.

#### Training

We train the MoTT network in a supervised way, using the cross-entropy (CE) loss to supervise the output of the classification head and the mean square error (MSE) loss to supervise the output of the regression head. The target of classification head output is a class index in the range [0, (*m* + 1)^*d*^) where (*m* + 1)^*d*^ is the number of hypothesis tracklets. The target of regression head output is the ground truth of the concatenation of normalized velocity and the existence flag.

#### Inference

In inference, we add a 1D max-pooling layer following the classification head to select the highest probability of the hypothesis tracklets with the same detection at frame *t* + 1 as the matching probabilities between the live tracklet and the candidate detection at frame *t* + 1. Then the (*m* + 1) predicted matching probabilities are normalized by softmax. The matching probabilities between the live tracklet and other detections besides the *m* + 1 candidate detections are set to zero.

### 2.3 Modeling discrete optimization problem

To find a one-to-one correspondence solution, we construct a discrete optimization formulation as (1), where *p*_*ij*_ is the predicted match probabilities between the live tracklet *i* and the detection *j*, and *a*_*ij*_ ∈ {0, 1} is the indicator variable. In particular, *j* = 0 represents the null detection.

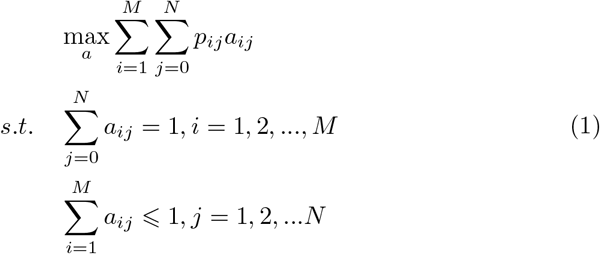

The objective function aims at maximizing the sum of matching probabilities under the constraints that each live tracklet is matched to only one detection (real or null), and each real detection is matched by at most one tracklet. This optimization problem is solved by using Gurobi (a solver for mathematical programming) [12] to obtain a one-to-one correspondence solution.

### 2.4 Track management

The one-to-one correspondence solution generally includes three situations. For each tracklet matched to a real detection, we add the matched real detection to the end of the live tracklet for updating. For each tracklet matched to a null detection, if the predicted existence probability is greater than a threshold *p* the predicted position is used to substitute for the null detection, else the live tracklet is terminated. In this way, the disconnected tracklets due to missing detections will be kept and be relinked when their detections emerge. For each detection that is not matched to any of the tracklets, a new live tracklet is initialized with this detection. After finishing particle linking on a whole movie, we remove the trajectories of length one, because they are considered false positive detections. See supplementary material for the details of track management.

## 3 Experimental Results

### Datasets

The performance of our method is evaluated on ISBI Particle Tracking Challenge datasets (ISBI PTC, http://bioimageanalysis.org/track/) [6], which consist of movies of biological particles of four subcellular structures: microtubule tips, vesicles, receptors, and viruses. These movies cover three different particle motion modes, four different SNR levels, three different particle density levels, and two different coordinate dimensions. For each movie in the training set, we use the first 70% frames for training and the last 30% frames for validation.

### Metrics

Metrics *α, β, JSC*_*θ*_, *JSC* are used to evaluate the method performance[6]. Metric *α* ∈ [0, 1] quantifies the matching degree of ground truth and estimated tracks, while *β* ∈ [0, *α*] is penalized by false positive tracks additionally compared to *α. JSC*_*θ*_ ∈ [0, 1] and *JSC* ∈ [0, 1] are the Jaccard similarity coefficient for entire tracks and track points, respectively. Higher values of the four metrics indicate better performance.

### Implementation details

In the following experiments, we set the length of live tracklets *Δt* + 1 = 7, the extension number *m* = 4, the depth of hypothesis tree *d* = 2, and the existence probability threshold *p* equals the mean of predicted existence probabilities of all live tracklets of current frame. See supplementary material for the ablation study on hyperparameters. We retrained the deepBlink network using simulated data generated by ISBI Challenge Track Generator. The MoTT model is implemented using PyTorch 1.8 and is trained on 1 NVIDIA GEFORCE RTX 2080 Ti with a batch size of 64 and an optimizer of Adam with an initial learning rate *lr* = 10^−3^, as well as *betas* = (0.9, 0.98) and *eps* = 10^−9^.

### 3.1 Quantitative Performance

#### Comparison with the SOTA methods

We compared our single particle tracking method with other SOTA methods, and the quantitative results on the microtubule scenario are shown in Table 1. Generally, our method outperforms other methods. Example visualization of tracking results can be found in Fig. 1.

**Table 1.** Comparison with SOTA methods on microtubule movies of ISBI PTC datasets. Method 5, Method 1, and Method 2 are the overall top-three approaches in the 2012 ISBI Particle Tracking Challenge. See [6] for details of these three methods. “−” denotes that results are not reported in the papers. Bold represents the best performance. Trackpy [1], SORT [28], Bytetrack [31] and Ours use the same detections.

| Density | Method | SNR = 4 |  |  |  | SNR = 7 |  |  |  |
| --- | --- | --- | --- | --- | --- | --- | --- | --- | --- |
| | | $\alpha$ | $\beta$ | $JSC_\theta$ | $JSC$ | $\alpha$ | $\beta$ | $JSC_\theta$ | $JSC$ |
| Low | Method5 | 0.750 | 0.728 | 0.917 | 0.874 | 0.803 | 0.787 | 0.939 | 0.894 |
|  | Method1 | 0.541 | 0.495 | 0.874 | 0.792 | 0.657 | 0.621 | 0.902 | 0.837 |
|  | Method2 | 0.562 | 0.259 | 0.356 | 0.369 | 0.694 | 0.686 | 0.959 | **0.954** |
|  | PMMS [22] | — | — | — | — | — | — | — | — |
|  | DPT [26] | — | — | — | — | — | — | — | — |
|  | SEF-GF-DPHT [25] | 0.803 | 0.776 | 0.928 | **0.890** | 0.861 | 0.848 | **0.970** | 0.936 |
|  | DetNet-DPHT [21] | 0.811 | 0.788 | 0.915 | 0.884 | 0.870 | 0.852 | 0.945 | 0.936 |
|  | Trackpy [1] | 0.762 | 0.657 | 0.749 | 0.694 | 0.853 | 0.789 | 0.854 | 0.808 |
|  | SORT [28] | 0.661 | 0.612 | 0.844 | 0.658 | 0.708 | 0.664 | 0.851 | 0.692 |
|  | Bytetrack [31] | 0.800 | **0.793** | **0.955** | 0.840 | 0.801 | 0.792 | 0.955 | 0.813 |
|  | Ours | **0.835** | 0.772 | 0.823 | 0.839 | **0.904** | **0.870** | 0.932 | 0.896 |
| Med | Method5 | 0.460 | 0.402 | 0.696 | 0.523 | 0.511 | 0.450 | 0.739 | 0.558 |
|  | Method1 | 0.353 | 0.264 | 0.550 | 0.373 | 0.400 | 0.326 | 0.646 | 0.448 |
|  | Method2 | 0.465 | 0.225 | 0.363 | 0.341 | 0.564 | 0.535 | **0.847** | 0.763 |
|  | PMMS [22] | 0.440 | 0.390 | 0.700 | 0.580 | — | — | — | — |
|  | DPT [26] | 0.488 | 0.373 | 0.556 | 0.449 | — | — | — | — |
|  | SEF-GF-DPHT [25] | 0.655 | 0.618 | **0.839** | 0.723 | — | — | — | — |
|  | DetNet-DPHT [21] | — | — | — | — | — | — | — | — |
|  | Trackpy [1] | 0.535 | 0.432 | 0.667 | 0.459 | 0.563 | 0.469 | 0.713 | 0.486 |
|  | SORT [28] | 0.544 | 0.478 | 0.733 | 0.528 | 0.583 | 0.523 | 0.757 | 0.558 |
|  | Bytetrack [31] | 0.555 | 0.495 | 0.717 | 0.552 | 0.582 | 0.528 | 0.721 | 0.567 |
|  | Ours | **0.814** | **0.719** | 0.760 | **0.769** | **0.869** | **0.792** | 0.829 | **0.823** |
| High | Method5 | 0.314 | 0.264 | 0.602 | 0.371 | 0.343 | 0.279 | 0.613 | 0.378 |
|  | Method1 | 0.272 | 0.210 | 0.544 | 0.299 | 0.293 | 0.231 | 0.582 | 0.322 |
|  | Method2 | 0.396 | 0.194 | 0.361 | 0.306 | 0.465 | 0.427 | 0.754 | 0.627 |
|  | PMMS [22] | 0.350 | 0.300 | 0.630 | 0.460 | — | — | — | — |
|  | DPT [26] | 0.414 | 0.313 | 0.524 | 0.389 | — | — | — | — |
|  | SEF-GF-DPHT [25] | 0.548 | 0.501 | **0.758** | 0.605 | — | — | — | — |
|  | DetNet-DPHT [21] | — | — | — | — | — | — | — | — |
|  | Trackpy [1] | 0.410 | 0.311 | 0.603 | 0.340 | 0.410 | 0.315 | 0.622 | 0.335 |
|  | SORT [28] | 0.432 | 0.354 | 0.645 | 0.407 | 0.465 | 0.390 | 0.664 | 0.436 |
|  | Bytetrack [31] | 0.385 | 0.313 | 0.558 | 0.377 | 0.425 | 0.354 | 0.593 | 0.407 |
|  | Ours | **0.732** | **0.611** | 0.660 | **0.659** | **0.814** | **0.718** | **0.759** | **0.753** |

#### Comparison under the same ground truth detections

Under the ground truth detections, we compare our particle linking method with LAP [14] and KF (Kalman filter) [15]. The results in Table 2 show that our method generally outperforms other methods in both medium-density and high-density cases.

**Table 2.** Comparison using the same ground truth detections on the microtubule, vesicle, and receptor scenarios.

|  |  | Microtubule |  |  | Vesicle |  |  | Receptor |  |  |
| --- | --- | --- | --- | --- | --- | --- | --- | --- | --- | --- |
| Density Method | | $\alpha$ | $\beta$ | $JSC_\theta$ | $\alpha$ | $\beta$ | $JSC_\theta$ | $\alpha$ | $\beta$ | $JSC_\theta$ |
| Low | LAP [14] | 0.850 | 0.852 | 0.923 | **0.953** | **0.947** | **0.979** | 0.940 | 0.931 | 0.962 |
|  | KF [15] | 0.972 | 0.962 | 0.971 | 0.937 | 0.924 | 0.959 | **0.964** | **0.955** | **0.972** |
|  | Ours | **0.988** | **0.985** | **0.993** | 0.926 | 0.891 | 0.925 | 0.949 | 0.921 | 0.943 |
| Med | LAP [14] | 0.486 | 0.394 | 0.662 | 0.753 | 0.703 | 0.704 | 0.742 | 0.686 | 0.826 |
|  | KF [15] | 0.827 | 0.798 | 0.859 | 0.673 | 0.609 | 0.787 | 0.824 | 0.794 | 0.867 |
|  | Ours | **0.992** | **0.987** | **0.992** | **0.800** | **0.733** | **0.874** | **0.930** | **0.894** | **0.935** |
| High | LAP [14] | 0.305 | 0.215 | 0.486 | 0.568 | 0.490 | 0.515 | 0.557 | 0.471 | 0.666 |
|  | KF [15] | 0.679 | 0.616 | 0.735 | 0.477 | 0.389 | 0.643 | 0.658 | 0.591 | 0.724 |
|  | Ours | **0.987** | **0.980** | **0.988** | **0.652** | **0.544** | **0.748** | **0.903** | **0.851** | **0.910** |

#### Effectiveness for all scenarios

We perform our particle linking method using ground truth detections on the four scenarios with three density levels in the ISBI PTC dataset. The results (see the supplementary material) demonstrate the effectiveness of our method for both 2D and 3D single particle tracking.

### 3.2 Robustness analysis

There are false positives (FPs) and false negatives (FNs) in actual detection results. Early study shows that FNs affect performance more than FPs [24]. We evaluated the robustness of our method under different FN levels. The receptor particle with medium density is used in this experiment. We randomly drop 5%, 10%, 15%, 20%, 30%, 40%, 50% detections from ground truth detections. As Fig. 4 shows, the tracking performance with the relinking strategy is better than that without the relinking strategy under different FN levels. Therefore, the proposed relinking strategy alleviates the impact of missed detections and enhances the robustness of our tracking method.

**Fig. 4.**
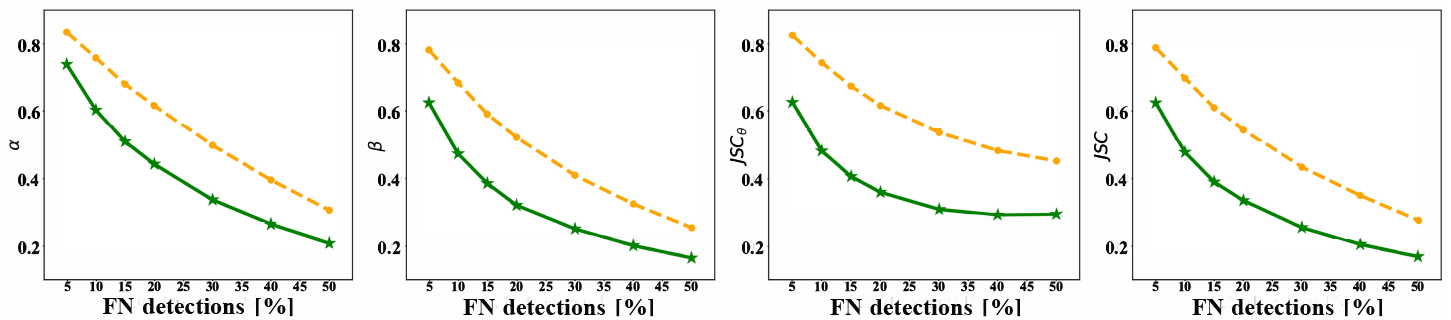
Robustness analysis under different levels of FN detection. The performance with the relinking strategy (orange) is better than that without the relinking strategy (green) under different FN levels.

## 4 Conclusion

In this paper, we proposed a novel Transformer-based method for single particle tracking in fluorescence microscopy images. We exploited the attention mechanism to model complex particle behaviors from past and hypothetical future tracklets. We designed a relinking strategy to alleviate the impact of missed detections due to e.g., low SNRs, and to enhance the robustness of our tracking method. Our experimental results show that our method is effective for all subcellular structures of ISBI Particle Tracking Challenge datasets, which cover different motion modes and different density levels. And our method achieves state-of-the-art performance on the microtubule movies of ISBI PTC datasets. In the future, we will test our method on other live cell fluorescence microscopy image sequences.

## Acknowledgements

This work was supported in part by the Natural Science Foundation of China (grants 31971289, 91954201) and the Strategic Priority Research Program of the Chinese Academy of Sciences (grant XDB37040402).

