## Supplementary material for "A Motion Transformer for Single Particle Tracking in Fluorescence Microscopy Images"

Table 1. Effectiveness for all scenarios in ISBI Particle Tracking Challenge [1].

| Object | Density | $\alpha$ | $\beta$ | $JSC_\theta$ | $JSC$ |
|---|---|---|---|---|---|
| Microtubules (directed 2D) | Low | 0.978 | 0.970 | 0.920 | 0.970 |
| | Middle | 0.994 | 0.991 | 0.978 | 0.991 |
| | High | 0.990 | 0.986 | 0.980 | 0.986 |
| Vesicles (brownian 2D) | Low | 0.903 | 0.865 | 0.904 | 0.856 |
| | Middle | 0.794 | 0.742 | 0.882 | 0.731 |
| | High | 0.709 | 0.637 | 0.822 | 0.638 |
| Receptors (switching 2D) | Low | 0.973 | 0.967 | 0.967 | 0.964 |
| | Middle | 0.952 | 0.939 | 0.949 | 0.939 |
| | High | 0.928 | 0.897 | 0.918 | 0.896 |
| Viruses (switching 3D) | Low | 0.999 | 0.999 | 0.997 | 0.999 |
| | Middle | 0.950 | 0.911 | 0.890 | 0.911 |
| | High | 0.951 | 0.916 | 0.901 | 0.916 |

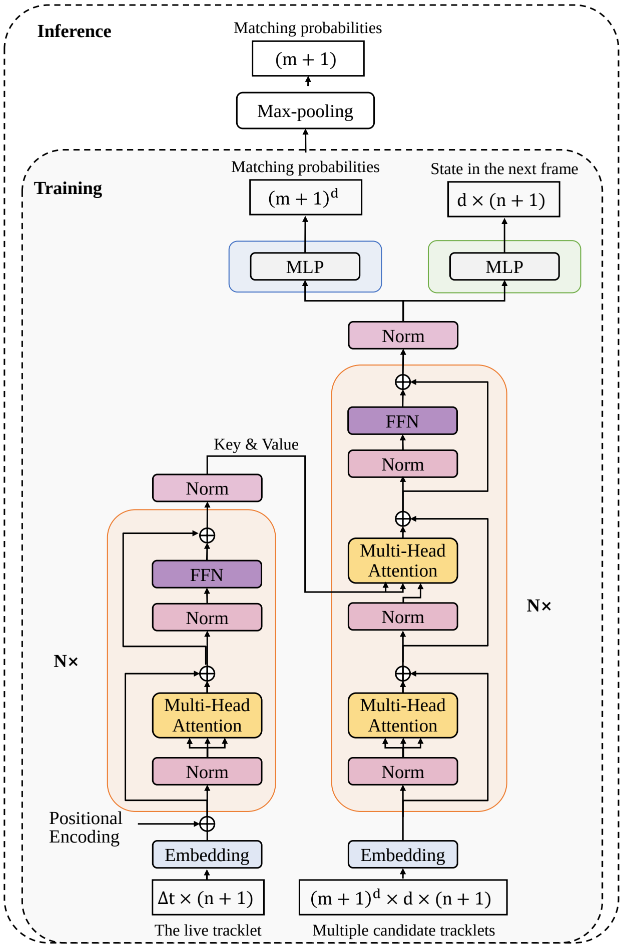


Figure 1. The detailed structure of the MoTT network. FFN: Feed-Forward Networks, MLP: Multilayer Perceptron.

**Algorithm 1** Management of tracks

**Input:** Live tracklet set $T_{live}$, result tracks set $T_{result}$, the solution from optimization problem, the threshold of existence probabilities $threshold$, predicted extension probabilities and predicted position from MoTT.

**Output:** Updated live tracklet set $T_{live}$ and result tracks set $T_{result}$.

**for** $T_i$ in $T_{live}$ **do**
  $Solution_i \leftarrow$ the matching solution of $T_i$
  **if** $Solution_i$ is a real detection **then**
    $T_i \leftarrow T_i.append(Solution_i)$
  **else**
    $p_i \leftarrow$ the predicted existence probability of $T_i$
    **if** $p_i > threshold$ **then**
      $pos_i \leftarrow$ the predicted next position of $T_i$
      $T_i \leftarrow T_i.append(pos_i)$
    **else**
      $T_{result} \leftarrow T_{result} \bigcup \{T_i\}$
      $T_{live} \leftarrow T_{live} \setminus \{T_i\}$
    **end if**
  **end if**
**end for**
**for** $D_j$ in unlinked detections of next frame **do**
  $T_{live} \leftarrow T_{live} \bigcup \{D_j\}$
**end for**
**return** $T_{live}, T_{result}$

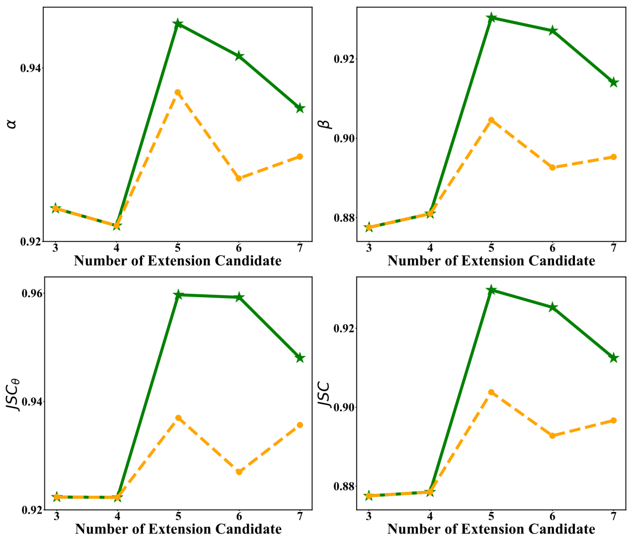


Figure 2. Ablation study about the number of extension candidates. There are two different network settings, one (green line) is {layer number = 1, head number = 6, k/v dimension = 96}, another (orange line) is {layer number = 2, head number = 2, k/v dimension = 64}. With five extension candidates, our method performs best.

**Algorithm 2** Preprocessing of tracklets

**Input:** One live tracklet $S$ and its candidate tracklet set $T_{cand}$; the constant length of live trackelts $(\Delta t + 1)$, the extension depth of hypothesis trees $d$; the number of hypothesis tracklets $m + 1$.

**Output:** The preprocessed tracklet $S_{pre}$ and candidate tracklets $T_{cand-pre}$.

/* Preprocessing of the live tracklet */
L $\leftarrow$ the length of S
**if** L $\leq (\Delta t + 1)$ **then**
    /* Padding with $\mathbf{-1}$ to make S length $\Delta t + 1$. */
    $S_c \leftarrow \{\mathbf{-1}, ..., \mathbf{-1}, D_S^{t-(L-1)}, ..., D_S^{t-1}, D_S^t\}$
**else**
    /* Clipping to make S length $\Delta t + 1$. */
    $S_c \leftarrow \{D_S^{t-\Delta t}, ..., D_S^{t-1}, D_S^t\}$
**end if**
**for** $D^i$ in $S_c \setminus \{D_S^t\}$ **do**
    **if** $D^i \neq \mathbf{-1}$ **then**
        $Shiftlive(i) \leftarrow D^{(i+1)} - D^i$
        $Flaglive(i) \leftarrow \mathbf{1}$
    **else**
        $Shiftlive(i) \leftarrow \mathbf{0}$
        $Flaglive(i) \leftarrow \mathbf{0}$
    **end if**
**end for**
$S_{pre} \leftarrow concatenate(Shiftlive, Flaglive)$
/* Preprocessing of multiple candidate tracklets*/
**for** $H_k$ in $T_{cand}$ **do**
    $LastD \leftarrow D_S^t$
    **for** $D^j$ in $H_k$ **do**
        **if** $D^j \neq \mathbf{-1}$ **then**
            $Shiftcand(k, j) \leftarrow D^j - LastD$
            $Flagcand(k, j) \leftarrow \mathbf{1}$
            $LastD \leftarrow D^j$
        **else**
            $Shiftcand(m, j), Flagcand(m, j) \leftarrow \mathbf{0}$
        **end if**
    **end for**
**end for**
$T_{cand-pre} \leftarrow concatenate(Shiftcand, Flagcand)$
**return** $S_{pre}, T_{cand-pre}$

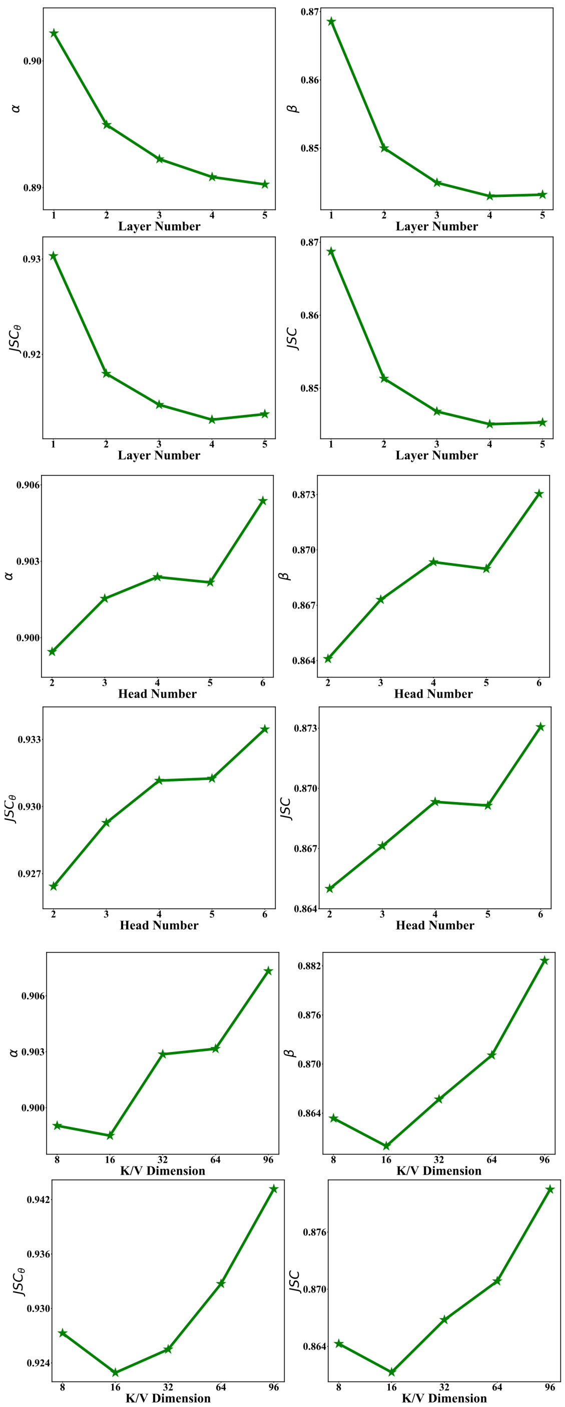


Figure 3. Ablation study of network hyperparameters. With the layer number increasing, the performance decreases. With the head number and key/value dimension increasing, the performance increases. With 1 Transformer layer, 6 heads, and 96 key/value dimension, the performance of our method performs best.
